# Sparse Stimulus Generation Improves Reverse Correlation Efficiency and Interpretability

**DOI:** 10.64898/2026.03.24.714012

**Authors:** Jane A. Gargano, Abigail Rice, Divya A. Chari, Benjamin Parrell, Adam C. Lammert

## Abstract

Reverse correlation is a widely-used and well-established method for probing latent perceptual representations in which subjects render subjective preference responses to ambiguous stimuli. Stimuli are purposefully designed to have no direct relationship with the target representation (e.g., they are randomly-generated), a property which makes each individual stimulus minimally informative toward reconstructing the target, and often difficult to interpret for subjects. As a result, a large number of stimulus-response pairs must be gathered from a given subject in order for reconstructions to be of sufficient quality, making the task fatiguing. Recent work has demonstrated that the number of trials needed can be substantially reduced using a compressive sensing framework that incorporates the assumption that the target representation can be sparsely represented in some basis into the reconstruction process. Here, we introduce an alternative method that incorporates the sparsity assumption directly into stimulus generation, which holds promise not only for improving efficiency, but also for improving the interpretability of stimuli from subject’s perspective. We develop this new method as a mathematical variation of the compressive sensing approach, before conducting one simulation study and two human subjects experiments to assess the benefits of this method to reconstruction quality, sample size efficiency, and subjective interpretability. Results show that sparse stimulus generation improves all three of these areas relative to conventional reverse correlation approaches, and also relative to compressive sensing in most conditions.

---

Reverse correlation is a data-driven approach to uncovering hidden representations, where responses are elicited from subjects following the presentation of stimuli with random properties. For example, in recent applications of reverse correlation to tinnitus characterization by Barnett et al. (2024), sounds are generated from spectra with random power levels at each frequency. Subjects are then asked to make subjective responses about whether they perceived the presence of the tinnitus sound they experience in the stimuli. A representation of the target percept can then be estimated that optimally explains the pattern of responses using one of several statistical procedures applied to these stimulus-response data. Statistical procedures range from restricted forms of linear regression (e.g., Gosselin & Schyns, 2003) to advanced signal reconstruction approaches (e.g., Roop, 2022; Roop et al, 2024). Reverse correlation has been widely used across a range of domains, including probing sensory sensitivity (Ahumada Jr & Lovell, 1971; De Boer & Kuyper, 1968), neural tuning (e.g., Ringach & Shapley, 2004; Nishimoto et al., 2006), cognitive representations (e.g., Ahumada Jr & Lovell, 1971; Gosselin & Schyns, 2001; Jäkel et al., 2009; Neri & Levi, 2006; Smith et al., 2012; Varnet et al., 2013a, 2013b) and psychological categories (e.g., Brinkman et al., 2017; Mangini & Biederman, 2004; Moon et al., 2020; Ponsot et al., 2018). Moreover, the method is mathematically related to “white noise” approach to system identification, which has been used for characterizing physiological (Marmarelis & Marmarelis, 1978) and engineered systems (Volterra, 1930; Wiener, 1958; Ljung, 1999).

In reverse correlation, the number of trials collected is a critical factor that determines the quality of the final reconstructions. Each stimulus provides only a small amount of information about the target, and must be aggregated with other stimuli and their respective responses in order to be useful. In practice, a large number trials are required to obtain a reconstruction of reasonable quality. For example, one widely-cited study using human face targets (Smith et al., 2012) had subjects complete 10,500 trials, a number which is not unusually large for such experiments, and which has been greatly exceeded in some studies (e.g., Gosselin and Schyns, 2003). This inefficiency limits the scope of analysis and the generality of conclusions that are possible given the time and effort involved in collecting sufficient data.

Moreover, the large number of required trials is likely to cause fatigue, which may affect participant responses. Indeed, participant fatigue is an acknowledged concern in reverse correlation studies (Verspui, 2018; Kevane and Koopmann-Holm, 2021; Varnet & Lorenzi, 2022). Many reverse correlation studies in the literature implicitly acknowledge the potential for fatigue in their study design, for example by dividing the large number of trials into blocks. Additionally, some studies have explicitly attributed portions of their study design to concerns over lack of participant motivation, for example by identifying and eliminating unmotivated participants from the data analysis (van Driel, 2017), or by incorporating estimates of noise due to attentional lapses into calculation of statistical significances (Dai and Michayl, 2010).

In reverse correlation, it is critical that the stimuli are richly varying within the dimensions of interest, to fully cover the space of perceptual features in a statistical sense. Variation is provided, in the vast majority of studies, by having a prominent, random component to the generation of stimuli. The resulting stimuli are considered ambiguous, which is ultimately concerning because of the high potential to cause confusion for subjects, stemming from the difficulty of perceiving and interpreting stimuli that have little in common with the target. Such confusion and the resulting cognitive effort may compound to fatigue caused by large numbers. Critically, this cumulative fatigue is likely to amplify the trial-wise inefficiency of reverse correlation by increasing noise in responses as a result of loss of attention and engagement, in the end negatively affecting overall reconstruction quality.

The presence and precise effect of subject confusion over ambiguous stimuli is understudied and little-represented in the literature. Anecdotally, subjects who participate in reverse correlation experiments in our lab frequently complain that the stimuli have little in common with their expectations relative to the target, and that the task is confusing as a result. In extreme situations, subjects have been known to find no acceptable stimuli at all, and simply respond in the negative to all stimuli, resulting in data that cannot be meaningfully analyzed. The ambiguous nature of typical stimuli used in reverse correlation also creates the need to carefully tailor subject instructions, as these have a profound impact on subject responses (Gezae et al. 2025). Even with careful instructions, however, participant confusion about ambiguous stimuli can be seen in low positive response rates. For example, Gosselin and Schyns (2003) instructed subjects that approximately 50% of stimuli should represent positive cases, but still observed positive responses as low as 11%.

One approach to mitigating subject confusion and fatigue, which can also improve the efficiency of reverse correlation, is to construct stimuli that are not strictly random, but are related to the target in some way. In situations where reasonable assumptions can be made about the nature of the target, those assumptions can be incorporated into the stimulus generation process. For example, it may be possible to generate stimuli for a study on face perception by overlaying noise on an image of a neutral face (e.g., Moon et al., 2020), or for a speech perception study by adding noise to recordings of natural speech (e.g., Varnet et al., 2019; Carrante et al., 2023). This approach would be expected to bias the reconstruction in ways delineated by the specific assumptions made and so must be used carefully, considering the acceptability of the bias introduced. Notably, it has been shown that stimuli can be conditioned statistically to improve efficiency while incorporating no prior knowledge about the target (Compton et al., 2023). Still, making some assumptions regarding the target, and therefore the form of the reconstruction, is a common approach with efficiency advantages.

One commonly made assumption regarding perceptual representations is sparsity - i.e., that the target can be well-represented by a small number of basis functions. Theoretically, sparsity is a sensible assumption because it is viewed as a fundamental organizing principle of human perceptual systems (Olshausen & Fields, 2004), and most real-world signals of interest are considered to be sparse or approximately sparse (Olshausen & Field, 1996; Srivastava et al., 2003; Bruckstein, Donoho & Elad, 2009; Marvasti et al., 2012). Practically, sparsity is a useful assumption because it reduces the number of free parameters that must be estimated as part of the reverse correlation reconstruction which, in addition to the number of trials collected, is the major factor influencing reconstruction quality. Reverse correlation is essentially a regression problem, regressing the subject responses onto the stimuli, to find a set of regression coefficients that can represent the target. In most applications of reverse correlation, the number of free parameters is equal simply to the dimensionality of the stimuli, as those dimensions correspond to regression coefficients, each one of which must have sufficient data support to be estimated to a reasonable quality. Assuming sparse representations allows for estimating a much smaller number of parameters, specifically those that are needed for the sparse representation. The approach of incorporating sparsity assumptions into reverse correlation was used in a logistic regression framework by Mineault, Berthelme & Pack (2009), and in a compressive sensing framework by Roop et al. (2024), both of which showed the potential for a substantial reduction in the number of required trials needed to achieve a given level of reconstruction quality compared to the standard regression approach.

Prior studies that incorporated the sparsity assumption into reverse correlation did so retrospectively - i.e., as part of the reconstruction procedure, after data had been collected. While this approach leads to efficiency gains and thus can reduce the number of trials needed, it does nothing to lessen the inherent difficulty of the task for subjects, and to mitigate the resulting fatigue and confusion that results from highly ambiguous stimuli in conventional reverse correlation experiments. The goal of this work is therefore to develop and assess a *prospective* approach to incorporating sparsity into reverse correlation. We call this new approach *sparse stimulus generation*. Using the compressive sensing framework described by Roop et al. (2024) as a starting point, we first develop the mathematical foundation of sparse stimulus generation. We then present evidence that sparse stimulus generation provides substantial efficiency advantages over standard approach to reverse correlation. First, we show results from simulation studies which allow us to systematically assess the advantages of the method and subsequently report on two studies with human participants which corroborate the effects expected from simulations and the mathematical formulation. Finally, we contrast the advantages of this approach relative to compressive sensing and highlight domains in which we believe sparse stimulus generation might be most useful.

## Formulation

To develop the present approach, we begin with the standard model of subject responses in reverse correlation, which has the form:

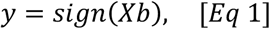

where *X* is a *n*-by-*m* stimulus matrix, *n* is the number of trials, and *m* is the dimensionality of the stimulus. The *m*-by-1 vector *b* is the signal of interest. The *n*-by-1 vector *y* contains yes/no responses coded as +1/-1, respectively, with use of the use *sign*(·) function.

One ubiquitous idea in the study of perception is that complex sensory stimuli are efficiently encoded using sets of features - either established or learned - in the form of salient stimulus patterns (Fairhall, 2014). For example, much effort has been dedicated to uncovering and deciphering spatial receptive fields in visual pathways (Movshon, Thompson & Tolhurst, 1978; DeAngelis, Ohzawa & Freeman, 1995), and spectro-temporal receptive fields in auditory pathways (Aertsen & Johannesma, 1981; David, Mesgarani & Shamma, 2007). Moreover, it is widely believed that these features are combined hierarchically to form increasingly abstract representations, and that higher-level representations draw sparsely from lower-level features (Olshausen & Field, 1996). Technological fields commonly use a parallel idea, that signals can be well-represented using some *basis* (e.g., Ahmed, Natarajan & Rao, 2006; Lee, 1996; Mermelstein, 1976; Turk & Pentland, 1991) and that sparse (Donoho, 2006; Baraniuk, 2007; Foucart & Rauhut, 2013) and/or hierarchical (Ng, 2011) representations are useful, owing to the widely-held belief that natural signals often have a sparse and/or hierarchical structure.

The notion of basis representations can be incorporated into reverse correlation in a variety of ways. For example, Roop et al. (2024) demonstrated that the efficiency of a reverse correlation experiment (i.e., the number of trials required to achieve an acceptable level of reconstruction accuracy) can be improved with the use of the signal processing technique compressive sensing. Compressive sensing begins with the assumption that the signal of interest (*b*) can be represented in some basis *Ψ* as follows:

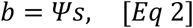

where *Ψ* is a *m*-by-*m* basis matrix, with basis vectors down the columns, *s* is a *m*-by-1 vector of coefficients.

The fundamental compressive sensing formula is:

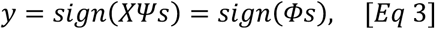

where *Φ* is a *n*-by-*m* matrix, sometimes called the *compressive sensing matrix*. Compressive sensing further assumes that the vector s will be sparse - i.e., having only *p* non-zero entries, where *p* < *m* - meaning that only a relatively small number of basis vectors from *Ψ* are needed to well-represent *b*.

In compressive sensing, the goal is to find the vector *s*, using some basis *Ψ*, a constructed stimulus matrix *X*, and a series of observed samples *y*. Under the assumption that *s* is sparse, it is possible to construct *X* such that *s* can be accurately estimated when *n* ≅ *p*, and such that *n* ≪ *m*.

In reverse correlation, an estimate of the signal of interest, *b̂*, is typically reconstructed as:

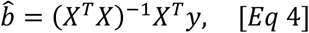

Using compressive sensing, Roop et al. (2024) showed that *b*^<^ can be estimated more efficiently as

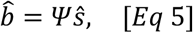

where *ŝ* = *γ*^−1^*P*(*m*^−1^*Φ^T^y*), *P* is a soft thresholding function, and *γ* = ‖*P*(*m*^−1^*Φ^T^y*)‖ is a normalization factor.

*X* is commonly constructed using random values, which allows estimation of arbitrary signals in reverse correlation, as well as statistical guarantees about efficiency improvements in compressive sensing. Indeed, this convergence is one reason why reverse correlation and compressive sensing are seen as compatible (Roop et al., 2024). While the most common approach to constructing stimuli in reverse correlation is to generate random noise in the dimensions that naturally compose the stimuli (e.g., pixel brightness for images, or frequency-domain power levels for sounds) the importance of bases, described above, points to an alternative approach in which random noise is generated instead in the dimensions defined by perceptual features or basis functions.

In the present formulation, as in compressive sensing, we begin with the assumption that *b* can be sparsely represented in some relevant basis, and that the basis is known prior to reconstruction. However, instead of incorporating that assumption into Eq 1 explicitly, as in compressive sensing (see Eqs 2 and 3), we instead support the assumption indirectly by generating stimuli within the space defined by a sparse selection of basis functions. Doing so further assumes that basis and the sparse selection of basis components is known somewhat earlier, prior to data collection, which differs from compressive sensing, wherein the sparse selection of basis components is estimated as part of the signal reconstruction process. If the sparse components are known prior to data collection, stimuli can be generated directly in the space defined by those components by randomly generating sparse vectors of coefficients.

Crucially, incorporating the sparsity assumption into the generation of stimuli addresses a fundamental problem with reverse correlation, which is that the random stimulus vectors traditionally used in reverse correlation have essentially zero correlation with the signal of interest with high probability, making the task difficult and error-prone for subjects. The proposed method is intended to benefit subjects, by making the typical stimulus more sensible to observe, allowing for more clear-cut and unambiguous response decisions.

Following this approach, we begin by assuming that *X* = *CSΨ^T^*, which allows us to reformulate Eq 1 as follows:

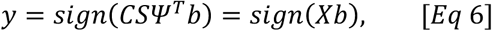

where *S* is an *m*-by-*m* diagonal matrix, where the *S_ii_* = 1 when |*s_i_*| > 0, and otherwise equal to 0. The matrix *C* is a *n*-by-*m* matrix of coefficients. Contrasting this formula with Eq 3 (compressive sensing), two main changes are apparent. Transposing *Ψ* makes it so that the coefficients being used to combine the basis vectors into a signal are in the rows of an *n*-by-*m* matrix *C*, whereas in Eq 3 those coefficients are in the vector *s*. Furthermore, constructing stimuli is no longer done by generating values in the matrix *X* directly, but rather by generating the values in the matrix *C*, which are then used to calculate the stimuli in combination with the basis vectors in *Ψ*. The matrix *S* specifies which basis vectors will be utilized as part of the sparse representation, but unlike the vector *s* from Eq 3, does not specify or influence the contribution of those basis vectors toward constructing the stimuli. Eq 6 also specifies that the signal of interest *b* is compared directly to the stimuli in *X*.

Using Eq 6, we begin to incorporate the assumption of sparsity. As with compressive sensing, we assume that some low-dimensional basis consisting of only *p* basis vectors can be used to represent a signal. However, whereas compressive sensing assumes that a low-dimensional representation of *b* is appropriate, we assume a low-dimensional representation of the stimuli - i.e., the rows of *X* - is appropriate and useful. That is, all but *p* coefficients in the rows of *C* will be unnecessary, because *p* columns of *Ψ* can be used to well-represent the stimuli. By determining the values in *C* randomly, then, these *p* vectors can be used to generate stimuli that populate the relevant *p*-dimensional subspace in which the signals of interest reside, rather than populating the larger *m*-dimensional space defined by the apparent dimensionality of that signal. Therefore, under the assumption of sparsity, it is entirely appropriate to rewrite Eq 6 more simply, without explicit reference to s as:

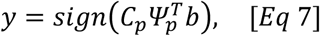

where *C_p_* is a *n*-by-*p* matrix and Ψ*_p_* is a *m*-by-*p* matrix, each containing only the columns of *C* and *Ψ* corresponding to |*s_i_*| > 0. Alternatively, this equation can be considered as abandoning the idea of sparsity altogether, and focusing attention on representations of the signal and stimuli that are simply low-dimensional – i.e., involving a small number of basis functions – and do not necessarily involve sparsity with respect to a larger set of basis functions. This formulation also makes it clear that *p* parameters are being estimated, which should be more efficient than estimating *m* parameters, as in Eq 1. In this new approach, an estimate of the signal of interest, *b*^<^, can be reconstructed using Eq 4, or equivalently using the following equation:

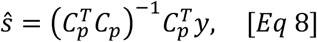

and finally,

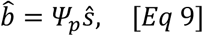

Note that the signal *b* will also be sparse in the same basis as the stimuli, because the stimuli are sparse, and because *b* will be a linear combination of stimulus vectors that are all sparse in the same basis. In brief, each stimulus is a linear combination of basis vectors, so a linear combination of stimuli is therefore a linear combination of linear combinations (i.e., a sum of sums). Therefore, this approach implicitly assumes what compressive sensing explicitly assumes, namely that *b* can be represented in some sparse basis.

Thus, for a given reverse correlation experiment, the idea of generating stimuli from a set of basis functions can be used in one of two ways: 1) if a large set of basis functions is domain-relevant, but a sparse subset of those functions is relevant to the task or target signal, Eq 6 allows the sparsity assumption to be included explicitly into the stimulus generation procedure, or 2) if a relatively low-dimensional set of basis functions (i.e., p < m) is relevant to the task or target signal outright, Eq 7 allows for that set to be used directly in stimulus generation.

A few studies have made use of closely-related approaches, sometimes under the name *dimensional noise*, often with the explicit goal of making reverse correlation experiments more efficient (Murray, 2011). One of the earliest applications of reverse correlation, an auditory study on tone detection, Ahumada & Lovell (1971) generated noise from a Fourier basis. In a study on stereoscopic disparity processing, Neri, Parker & Blakemore (1999) generated noise using a collection of spatially localized “dots” that were relatively large and few in number compared to image pixels. Levi & Klein (2002), investigating visual detection and position discrimination, generated noise using a parameterized sinusoidal basis. Li, Klein, & Levi (2006), in a further study on position acuity, generated noise in the form of parameterized spatial shape distortions. These studies have shown good levels of efficiency, likely owing to their use of the dimensional noise approach. Even so, there has been no systematic investigation of the benefits of this approach to efficiency.

Mangini and Biederman (2004) took the ideas of noise bases and sparsity one step further. In their visual study of face classification, they defined a basis of 4092 sinusoids over images composed of 16384 pixels, and noted efficiency gains following from this reduced dimensionality. During analysis, the data were used to identify a further reduced number of basis components - on the order or 1-5% of the original basis functions - that were nonetheless sufficient for an accurate but sparse representation of the target signal. Although not attempted by Mangini and Biederman (2004), these sparse components could have been used to generate stimuli for subsequent behavioral studies, in accordance with the present approach, to further improve efficiency by generating stimuli that are even more directly relevant to the target signal.

The literature is seemingly rife with examples of low-dimensional and/or sparse bases that would be compatible with our proposed approach. Prominent examples include bases visual perception of natural scenes (Olshausen & Field, 2004; Ringach & Shapley, 2004; Hunt, Dayan & Goodhill, 2013), perception of human faces (Turk and Pentland, 1991) and human voice (Kuhn et al., 1998), and speech perception at the cognitive (Schroeder, 1967; Mermelstein, 1967) and neurologic (Klein, Koenig & Koerding, 2003; David, Mesgarani & Shamma, 2008; Carlson, Ming & DeWeese, 2012) levels.

## Experimental Methods

One simulation study and one behavioral experiment were conducted, each with the aim of determining whether sparse stimulus generation has an impact on efficiency compared to conventional, nonsparse stimulus generation in reverse correlation experiments. In addition, a second behavioral experiment was conducted with the aim of assessing the subjective qualities of sparse stimuli. In the simulation study, we assessed reconstruction accuracy using a response model (i.e., Eq 1) across a range of stimulus sparsity levels and (simulated) numbers of trials. In the two behavioral studies, human participants completed a parallel experiment, in which responses were collected to either sparsely generated or conventional stimuli. In the simulation study and the first behavioral experiment, the quality of individual reconstructions and the overall effect of sparse stimulus generation on efficiency of reverse correlation was assessed by comparing reconstructions against a standard reference, and identifying the number of trials required to achieve a specific level of reconstruction quality. As a test case, we examine vowel perception, an area where reverse correlation has proved to be applicable (Brimijoin et al., 2013; Varnet et al., 2013b, Varnet et al., 2013b). Additionally, there is a well-known sparse basis within the domain for describing vocal tract shapes for vowels (Schroeder, 1967; Mermelstein, 1967), and that basis has a well-understood relationship with speech acoustics and, consequently, with speech perception (Liberman & Whalen, 2000; Galantucci, Fowler & Turvey, 2006; Lammert et al., 2025).

In speech production, the vocal folds near the glottis vibrate to produce sound waves at the beginning of the vocal tract (Stevens, 2000). As these sound waves move through the vocal tract, they are modified to create a unique sound. The vocal tract can be considered as having the effect of a set of band-pass filters (Johnson, 2011), emphasizing certain frequencies over others. Therefore, changing the shape of the vocal tract changes the sound produced. On the perceptual side, speech is generally thought to be perceived (at least quasi-) categorically (Johnson, 2011), such that listeners evaluate sounds they hear based on categories they have learned are important, tending to group sounds together that correspond to sounds frequently used in speech.

The current experiments are based on a previous study using reverse correlation to assess vowel perception (Lammert et al., 2025). In that work, human listeners were asked to listen to ambiguous auditory stimuli and provide responses as to whether a given stimulus matched a target vowel (i.e., “was the sound the vowel in *heed*?”). Subjects provided simple yes/no responses to this prompt. The auditory stimuli stemmed from random vocal tract shapes in a simulated vocal tract, which in turn came from random-generated coefficients of a cosine basis for describing vocal tract shapes (Schroeder, 1967; Mermelstein, 1967). Resonant sounds were synthesized from those vocal tract shapes and played to listeners. This task constitutes a reverse correlation experiment, which allows for typical reverse correlation reconstruction techniques to be applied, resulting in an estimated vocal tract shape corresponding to the target vowel. Although this application provides a useful testbed in which to examine the potential benefits of sparse stimulus generation given the strong evidence for sparse basis functions in vocal tract shapes, we note that the scientific and theoretical merit of this question is not critical to our aim of establishing the efficacy of the proposed method.

## Speech production basis

One appropriate basis for describing vocal tract area functions is a series of cosine functions, evaluated along the long axis of the vocal tract from the glottis to the lips (Mermelstein 1967). Using only six low-frequency cosines produces vocal tract shapes with low error compared with measured vowel shapes, and captures those basis functions that most directly relate to the three lowest formant frequencies (Schroeder, 1967). Therefore, using this basis as a low-dimensional representation of vocal tract area functions is closely related to the standard use of formant frequencies as a low-dimensional representation of speech acoustics. In addition, a cosine basis is a common representation of cognitive stimuli, which means it may represent an ideal testing basis in order to move towards applying sparse stimulus generation to other sparse domains (Concetta Morrone & Burr, 1988). At the same time, using the basis over a much wider range of frequencies – potentially up to p=m such frequencies – allows for variable accuracy, and degree of sparsity, of the representation.

The cosine basis can be stated formally as:

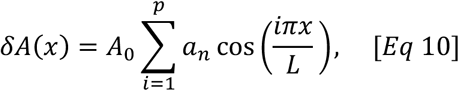

where *x* is the distance along the vocal tract, *L* is the overall length of the vocal tract, *i* is the spatial frequency of a single basis function and *a_i_* is the coefficient determining the contribution of that component. Given a set of coefficients *a*_1…*p*_, this formula can be used to calculate an area function as *A*(*x*) = *A*_0_ + *δA*(*x*). This formula is used to yield Ψ*_p_* is a *m*-by-*p* matrix, as in Eq 7, here containing values such that column *n* constitutes basis function *n*.

## Measured vocal tract shape

To facilitate the assessment of reconstruction quality in our vowel perception experiments, it is useful to have a standard vocal tract shape to compare reconstructions against. The standard shape used in this case comes from an MRI capture of the static vocal tract when making the vowel /i/ (as in “heed”), as reported by Story et al. (1996). Story captured a speakers’ midsagittal plane at 5 mm slice thickness, and analyzed the resulting image in order to measure the cross-sectional area of the vocal tract at 42 points along its length. This information was used to estimate the area function of the vocal tract, a concrete representation of the shape of the vocal tract during production of the vowel /i/, which can be directly compared to the reconstructions obtained from reverse correlation. We label this standard shape *b*, and call it the *target* signal in the present context. We note that using this target does assume that the vocal tract shape that best explains a listeners vowel perception behavior corresponds closely to the vocal tract shapes typically exhibited by speakers during production. Based on the results report by Lammert et al. (2025), in which a close correspondence between this target and reconstructed vocal tract shapes was observed, this assumption is a reasonable approximation.

Sparsified versions of this target signal derived from different numbers of basis functions, *p*, are shown in Figure 1. These renderings represent a reference for understanding the effects of sparsity on the target signal. To obtain these examples, the target signal was first projected onto Schroeder’s cosine basis, as 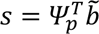, and coefficients *s* were subsequently used to calculate the target as *Ψ_p_*s. Figure 1 contains four graphed representations of the target signal – i.e., the measured /i/ vocal tract reported by Story et al. (1996) – at varying levels of sparsity: *p* = 42, *p* = 30, *p* = 18, and *p* = 6.

**Figure 1.**
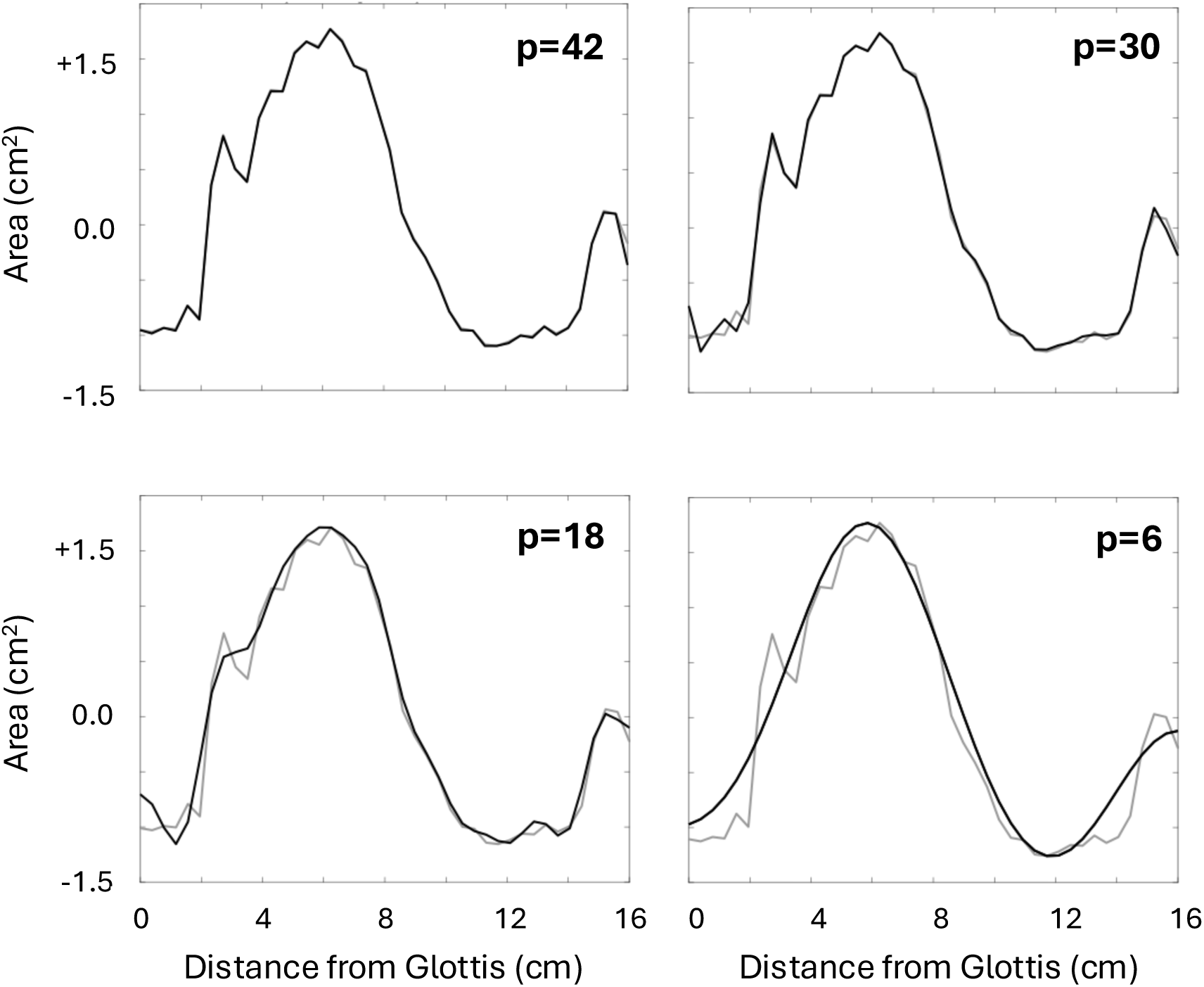
Representations of the /i/ vocal tract target shape at different levels of sparsity. The measured vocal tract shape is shown for reference (gray line). As the number of basis functions, *p*, goes from 42 down to 6, the representation appears simpler (e.g., smoother, less detailed). However, even when the value of *p* is quite low, the overall shape of the vocal tract is preserved, implying its important characteristics can be represented sparsely, i.e., using only a small subset of the full basis.

## Acoustic synthesis

To transform vocal tract shapes to the acoustic stimuli presented to participants, we used a common approach where the vocal tract is treated as a series of concatenated, cylindrical tubes (Kelly & Lochbaum, 1962; Iskarous, 2010; Lammert & Narayanan, 2015). In this case, assumptions are made that the vocal tract has a simplified geometry, no loss, and no radiation impedance. It is well-established that the reflection coefficients between adjacent tubes can be calculated using the formula:

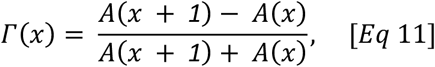

where *A*(*x*) is the cross-sectional area of the vocal tract at distance *x* from the glottis. The coefficients of a prediction filter polynomial can then be calculated from these reflection coefficients, the formant frequencies are then calculated as the roots of that polynomial. Bandwidths for each formant frequencies are calculated using the approximation due to Fant (1985). Finally, the calculated frequency and bandwidth of each formant are used to define the filter coefficients of a digital resonator which is used to filter a synthesized glottal signal as part of a filter cascade. The glottal signal was a constructed as a train of pulses occurring at 180Hz. Filters corresponding to the lowest three formants were used to filter the signal, producing the acoustic waveform used as stimuli.

## Simulation study

Simulations were conducted to assess the accuracy of the reverse correlation reconstructions, *b*^<^, as compared to a target signal, *b*, using sparse stimulus generation as a function of the number of basis functions used to generate stimuli, *p*, and the number of trials completed by the simulated subject, *n*. The amount of perceptual noise was assumed to be negligible. The number of trials considered ranged from a minimum of *m* up to a maximum of 4*m*, in increments of 4. The number of basis functions considered ranged from a minimum of 4 up to *m* = 42, in increments of 2. For each simulation, reconstruction quality was assessed by calculating Pearson’s *r* correlation coefficient between *b* and *b*^<^ as estimated in that simulation. For every choice of *n* and *p*, a total of 100 simulations were performed, and the reconstruction quality measure was averaged across those 100 simulations.

In simulating the sparse stimulus generation approach, responses were generated as in Eq 7, where *C_p_* was a *n*-by-*p* matrix of random coefficients uniformly distributed between -1 and +1, and Ψ*_p_* is a *m*-by-*p* matrix such that column *i* constitutes basis function *i* from Eq 10. Reconstructions, *b*^<^, in the sparse stimulus generation approach were obtained using Eq 4. This was contrasted with a conventional reverse correlation approach, here referred to as “nonsparse stimulus generation”, in which responses were generated as in Eq 1, where *X* was an *n*-by-*m* stimulus matrix with values uniformly distributed between -1 and +1. Reconstructions, *b*^<^, in this “nonsparse” approach were obtained from Eq 4, or alternatively using Eq 5 (i.e., using compressive sensing), where *Ψ* was an *m*-by-*m* matrix such that column *i* constitutes basis function *i* from Eq 10. The target signal, *b*, was taken to be the measured vocal tract shape reported by Story et al. (1996), as described above.

In the case of the current study, we modeled subject response direction on the basis of stimulus matrixes *X* containing vocal tract area functions *A*, rather than generating acoustic signals as in the behavioral studies below. Our assumption here is that listeners can essentially invert the acoustic signal to perceive the underlying vocal tract (Liberman & Whalen, 2000; Galantucci, Fowler & Turvey, 2006). Building on the subject response model from Eq 1, this concept can be stated more precisely with an augmented subject response model: *y* = *sign*I*F*^−1^I*F*(*A*)J*b*J, where the function *F*(·) represents the physical process by which vocal tract shapes generate an acoustic signature. In this interpretation, we do not know the function *F*(·), nor do we care much what it is in specific terms, but for the purposes of this simulation we assume that listeners have a cognitive process which inverts the function *F*(·), i.e., *F*^−1^(·), so that perceptual judgements may be made in articulatory terms by comparing stimuli against the internal representation *b*, which is a cognitive representation of the vocal tract shape associated with the target vowel. From these assumptions, specifically the fact that *F*^−1^I*F*(*A*)J = *A*, the subject response model can be rewritten as *y* = *sign*(*Ab*), which can be inverted as in Eq 4 to be written as: *b̂* = (*A^T^ A*)^−1^*A^T^y*.

## Human subjects experiment 1

Trials with human subjects were also conducted with the aim of reconstructing vocal tract shapes consistent with the vowel /i/ on the basis of both sparse stimuli and random stimuli, and assessing the quality of those reconstructions relative to the same measured vocal tract shape used as a target signal in the simulation study. For sparse stimuli, six basis functions were used in generating vocal tract area functions for the stimuli (*p* = 6), and these area functions were sampled at 32 points along the vocal tract (*m* = 32). In the nonsparse case, the area of the vocal tract was randomly determined at each of *m* = 32 points along the vocal tract without considering basis functions.

Sparse stimuli were generated as in Eq 7, where *C*_*_ was a 200-by-6 matrix of random coefficients uniformly distributed between -1 and +1, and Ψ_*_is a 32-by-6 matrix with columns representing the six lowest basis functions from Eq 10. Reconstructions, *b̂*, in the sparse stimulus generation approach were obtained using Eq 4. Conventional (i.e., “nonsparse”) stimuli were generated as in Eq 1, where *X* was an *n*-by-*m* stimulus matrix with values uniformly distributed between -1 and +1. Reconstructions, *b̂*, in this “nonsparse” approach were obtained both from Eq 4 (conventional reverse correlation) and using Eq 5 (using compressive sensing), where *Ψ* was an 32-by-32 matrix such that column *i* constitutes basis function *i* from Eq 10.

Three listeners with self-reported normal hearing were recruited for experiment 1. All procedures were approved by the UMASS IRB (H00024058). Subjects listened to stimuli through headphones in a quiet listening environment and were instructed to respond via keyboard press to each sound in response to the prompt: “did you hear the vowel ‘ee’ as in ‘heed’”; “yes” (f key) or “no” (j key). Subjects completed a total of 200 trials each with the sparse and nonsparse stimuli, presented in eight blocks of 50 trials of one condition type, presented in a random order. Reconstructions, *b̂*, were once again produced using Eq 4, and also using compressive sensing, as described in Eq 5.

Quality of the reconstruction was assessed, as in the simulation study, using Pearson’s *r* correlation coefficient between *b* and *b*<, after resampling *b* to m=32. To facilitate the assessment of reconstruction quality across a range of trial numbers, a bootstrap procedure was employed in which the data collected were resampled with replacement into random subsets (i.e., bootstrap samples) of stimulus-response pairs ranging in size from 40 to 200. For each bootstrap sample size, resampling was performed 30 times, and mean correlation values and 95% confidence intervals were calculated.

## Human subjects experiment 2

Here, we tested whether subjective fatigue and confusion with respect to stimuli were improved using sparse stimuli, as compared to conventionally generated stimuli. Six subjects with self-reported normal hearing completed experiment 2. The order of the blocks was randomized within and across subjects. All procedures were approved by the UMASS IRB (H00024058).

Subjects completed 100 reverse correlation trials each with sparse and nonsparse stimuli, presented in eight blocks of 25 trials. Stimulus generation and participant instructions were the same as for human subjects experiment 1. After each block of 25 trials, subjects were asked to agree or disagree with two statements about the tasks, using on a Likert scale ranging from 1 to 7 (1 = strongly agree, 4 = neither agree nor disagree, 7 = strongly disagree). The first statement, aimed at assessing task difficulty (and therefore the potential for subject fatigue), was: “some sounds were a lot like the target vowel”. The second statement, aimed at assessing subject confusion was: “when I answered yes, I felt confident”.

Though this purpose of this experiment was not to assess reconstruction quality, reconstructions were nevertheless obtained for these subjects, and the quality of reconstructions was assessed as in human subjects experiment 1. These reconstructions are shown in Figure 6, for the sake of completeness.

## Results

### Simulation study

Figure 2 shows the simulation study results. Subfigures show the estimation quality of simulated vocal tract reconstructions, in terms of Pearson’s *r* with the target signal *b*, as contours according to the number of basis functions (*p*) and the number of trials completed (*n*) used to generate the reconstruction. Each subfigure shows estimation quality corresponding to a different stimulus generation and reconstruction strategy – from left to right: nonsparse stimulus generation with conventional reverse correlation reconstruction (Eq 4), nonsparse stimulus generation with compressive sensing reconstruction (Eq 5), and sparse stimulus generation with conventional reconstruction (Eq 4). The presented quality measures represent the mean value over 100 repetitions of the simulation at corresponding values of *p* and *n*.

**Figure 2.**
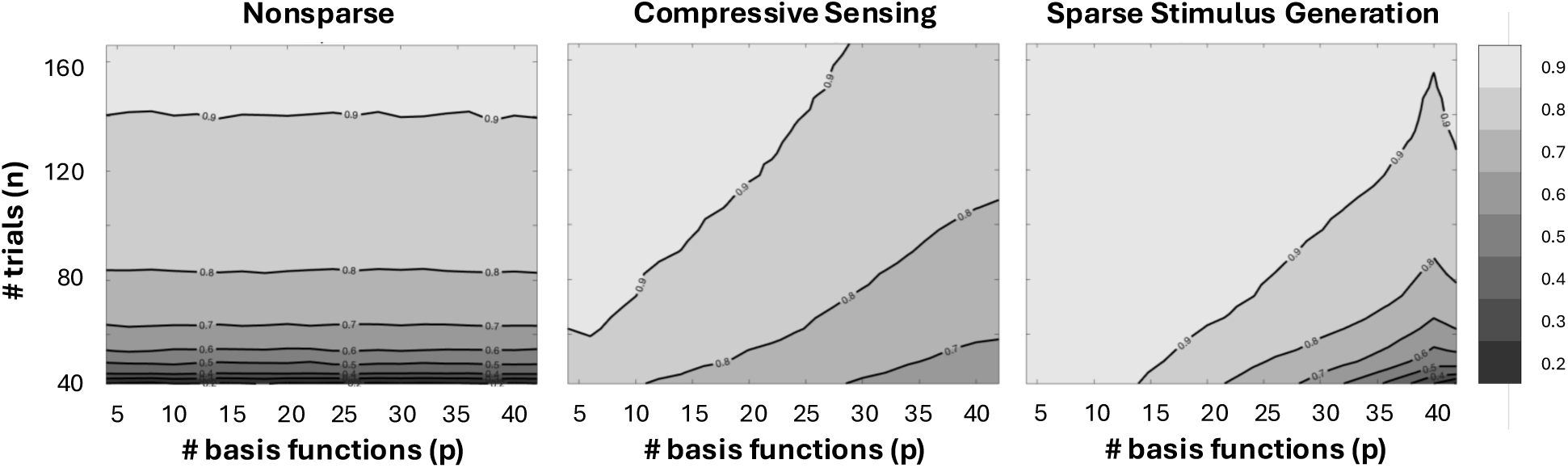
Simulation study results showing estimation quality relative to the target vocal tract shape in terms of Pearson’s r and as a function of number of basis functions (p) and number of trials completed (n). Subfigures show results using (left to right) nonsparse stimuli, compressive sensing, and sparse stimulus generation. The estimation quality values shown represent the mean value over 100 repetitions of the simulation at corresponding values of p and n. Results show that the sparse stimulus generation approach displays a higher reconstruction quality than the nonsparse approach for nearly any value of p and n.

In the simulation study results, the sparse stimulus generation approach displays a higher reconstruction quality than the nonsparse approach for nearly any value of *p* and *n*. For example, at sparsity *p* = 22, sparse stimulus generation requires approximately half the number of trials compared to nonsparse stimulus generation to reach accuracy above 0.9. This relative gap varies with sparsity, as the number of required trials to reach any particular accuracy decreases with increasing sparsity using the sparse stimuli approach (i.e., as *p* decreases), while the number of required trials using the nonsparse approach is (expectedly) unaffected by the value of *p*. When sparsity levels are very low (i.e., as *p* ≈ 42, the dimensionality of the original signal *b*), the performance of the sparse and nonsparse approaches is largely similar.

Comparing sparse stimulus generation versus the compressive sensing approach, it is evident that sparse stimulus generation consistently displays a higher reconstruction quality across a wide range of values *p* and *n*. For example, to reach accuracy above 0.9 at sparsity *p* = 14, compressive sensing requires more than double the number of trials compared to sparse stimuli. This relative gap varies only slightly with sparsity, as both compressive sensing and sparse stimulus generation appear similarly sensitive to the value of *p*, and in general approximately 75% more trials are required for compressive sensing than for sparse stimulus generation.

Figure 3 shows some example reconstructions of the target vocal tract shape taken from the simulation study. Subfigures show, across the columns, reconstructions for a fixed level of sparsity (*p* = 6) using, from left to right: nonsparse stimulus generation with conventional reverse correlation reconstruction (Eq 4), nonsparse stimulus generation with compressive sensing reconstruction (Eq 5), and sparse stimulus generation with conventional reconstruction (Eq 4). Subfigure rows represent reconstructions stemming from a very small (*n* = *m*) or rather large (*n* = 4*m*) number of trials.

**Figure 3.**
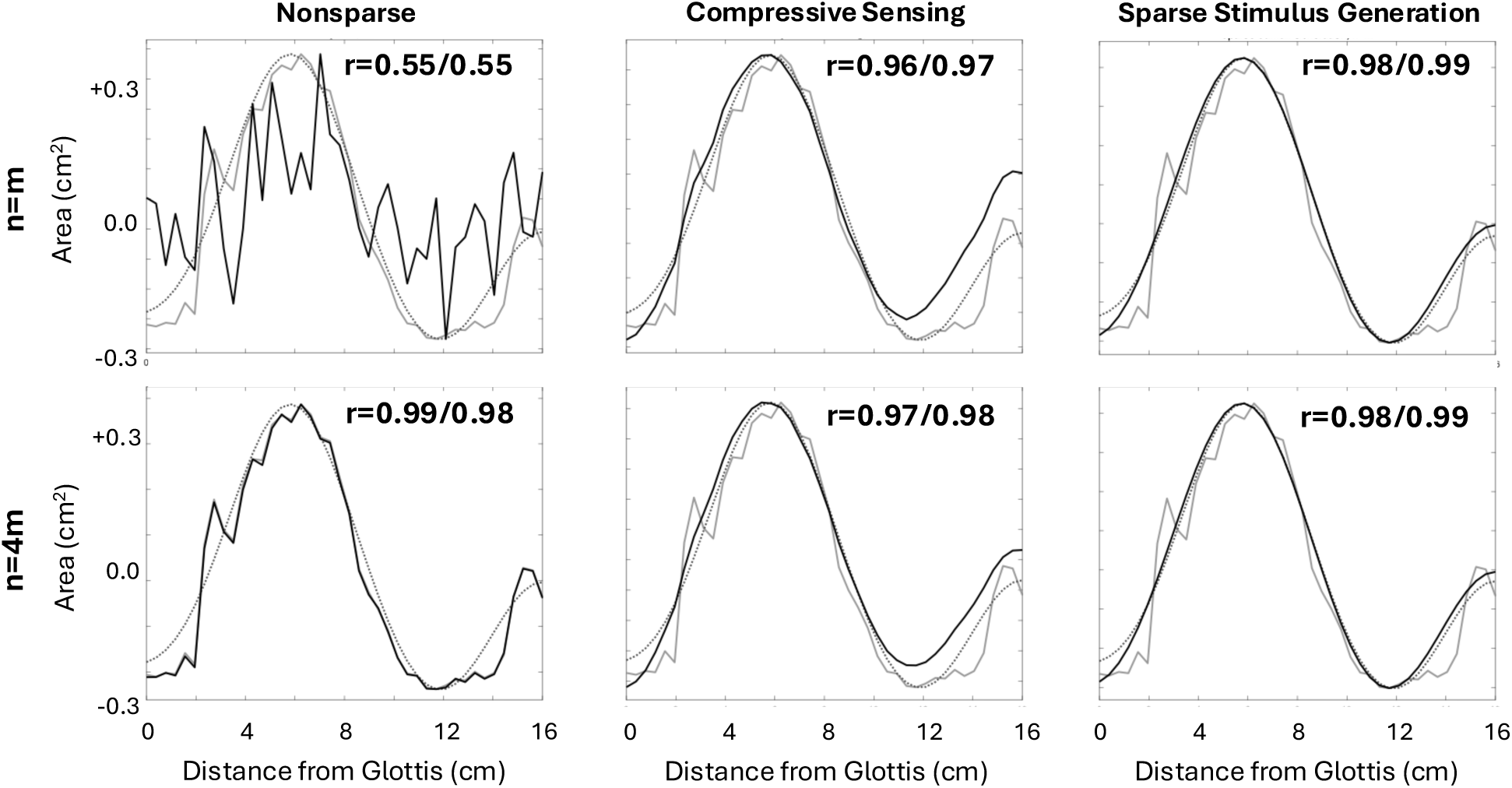
Example reconstructions from the simulation study of the target vocal tract shape. Subfigures show reconstructions for a fixed level of sparsity (p=6) using the nonsparse, compressive sensing and sparse stimulus generation approaches (columns, left to right) and for a small (n=m) and large (n=4m) number of trials (rows, top to bottom). Estimation quality relative to the target signal b (gray, solid line), and a sparsified (p=6; gray, dotted line) version of b (as in Figure 1) are shown, respectively.

Estimation quality in terms of Pearson’s *r* between the reconstruction and both the original target signal *b* (left of slash), as well as a sparsified (*p* = 6) version of *b* (right of slash).

In the example reconstructions (Figure 3), sparse stimulus generation reconstructions have correlation values that are above 0.98 in all cases. There was only moderate correlation (∼0.55) between the nonsparse reconstruction and the original reconstruction when using only m (=42) trials, but a large increase can be seen in the correlation between the nonsparse version at 4*m* trials as compared to *m* trials, from around 0.55 to above 0.98. The compressive sensing representations had similar albeit slightly lower correlation values to the sparse stimulus generation correlation values. Comparing the reconstructions obtained from the nonsparse approach to those obtained from sparse stimulus generation and compressive sensing in the large number of trials (*n* = 4*m*) condition, all reconstructions exhibit a very high correlation value with the target signal. Yet, more of the fine details in the target signal are apparent in the nonsparse reconstruction, by visual comparison with the target signal in Figure 1.

### Human subjects experiments

Figure 4 shows the results of the human subjects experiment 1. Subfigures show the simulation quality, in terms of mean Pearson’s *r* value across bootstrap iterations, as a function of the number of trials (*n*). The three lines in each plot indicate this reconstruction quality for nonsparse stimulus generation with conventional reverse correlation reconstruction (Eq 4), nonsparse stimulus generation with compressive sensing reconstruction (Eq 5), and sparse stimulus generation with conventional reconstruction (Eq 4).

**Figure 4.**
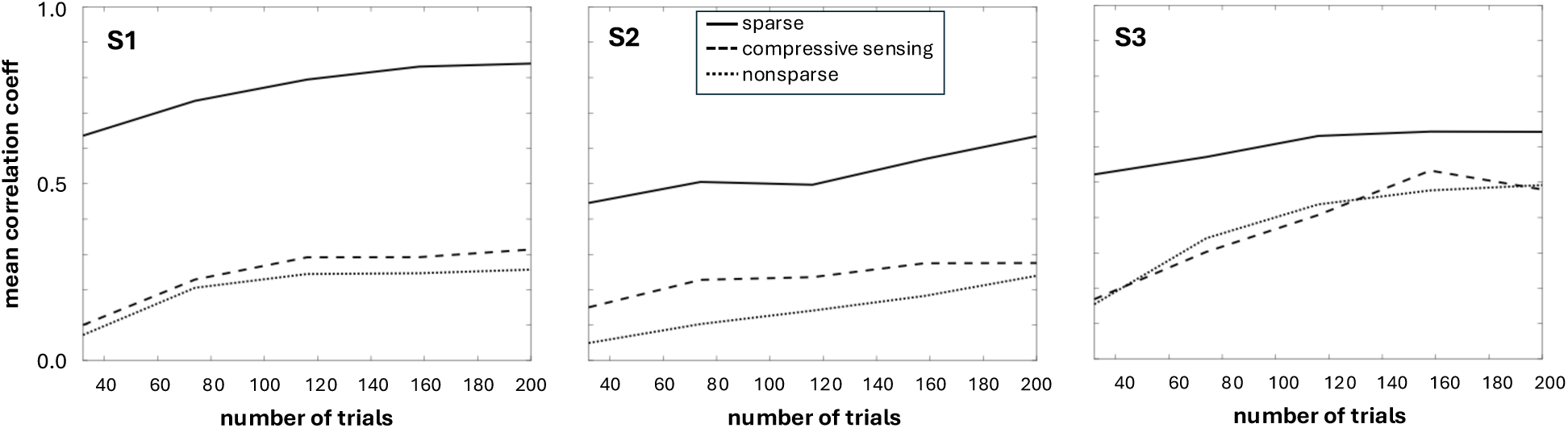
Human subjects results showing reconstruction quality relative to the target vocal tract shape in terms of Pearson’s r and as a function of number of the number of trials (n). Subfigures show the reconstruction quality for different subjects, with the three lines in each plot indicating reconstruction quality using nonsparse stimulus generation, compressive sensing, and sparse stimulus generation.

Figure 5 shows reconstructions of the target vocal tract shape taken from experiment 1.

**Figure 5.**
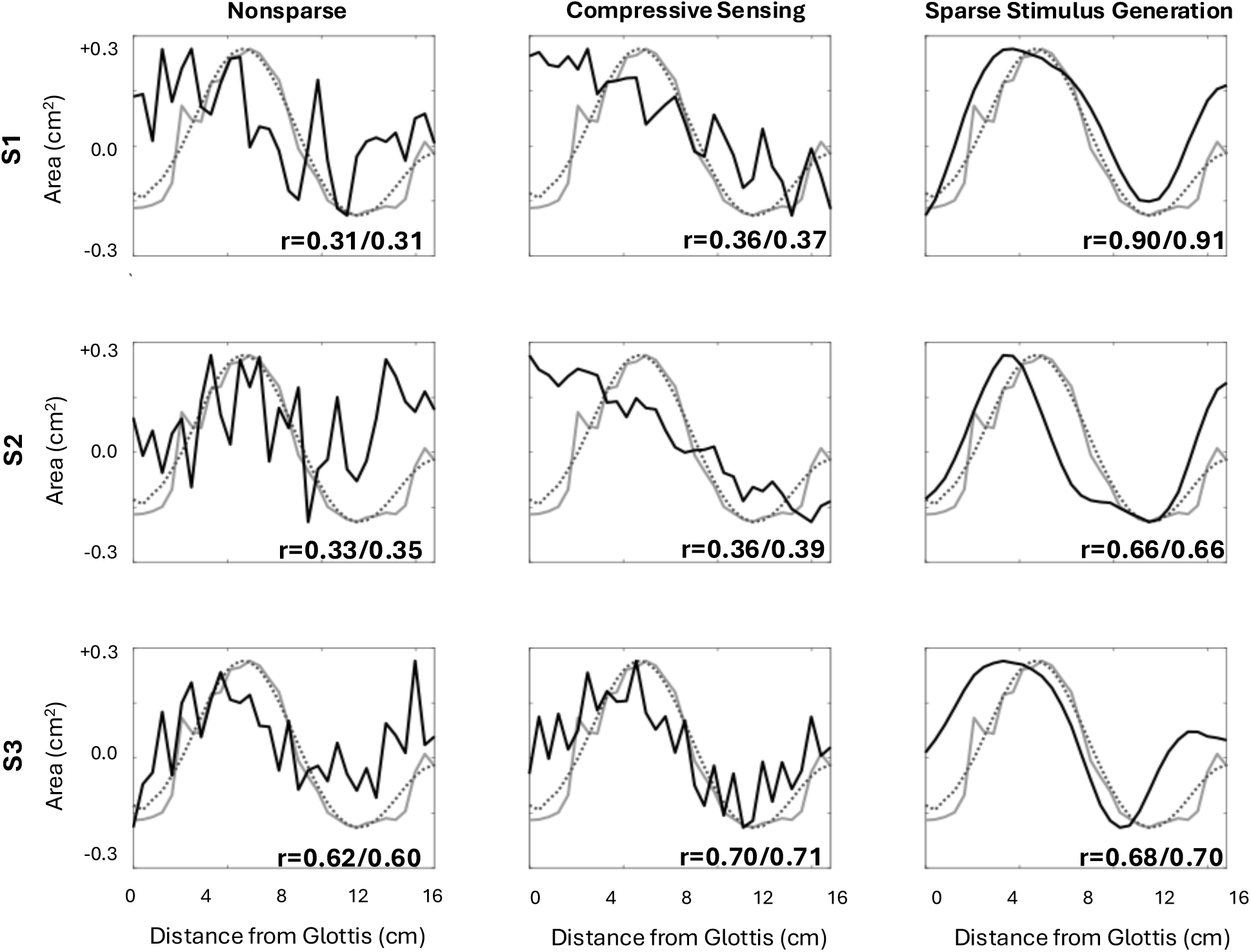
Example reconstructions from human subjects. experiment 1 of the target vocal tract shape. Subfigures show reconstructions for a fixed level of sparsity (p=6) using the nonsparse, compressive sensing and sparse stimulus generation approaches (columns, left to right) and for a fixed number of trials (n=200) for each subject (rows). Estimation quality relative to the target signal b (gray, solid line), and a sparsified (p=6; gray, dotted line) version of b (as in Figure 1) are shown, respectively (target signal was resampled to m=32).

**Figure 6.**
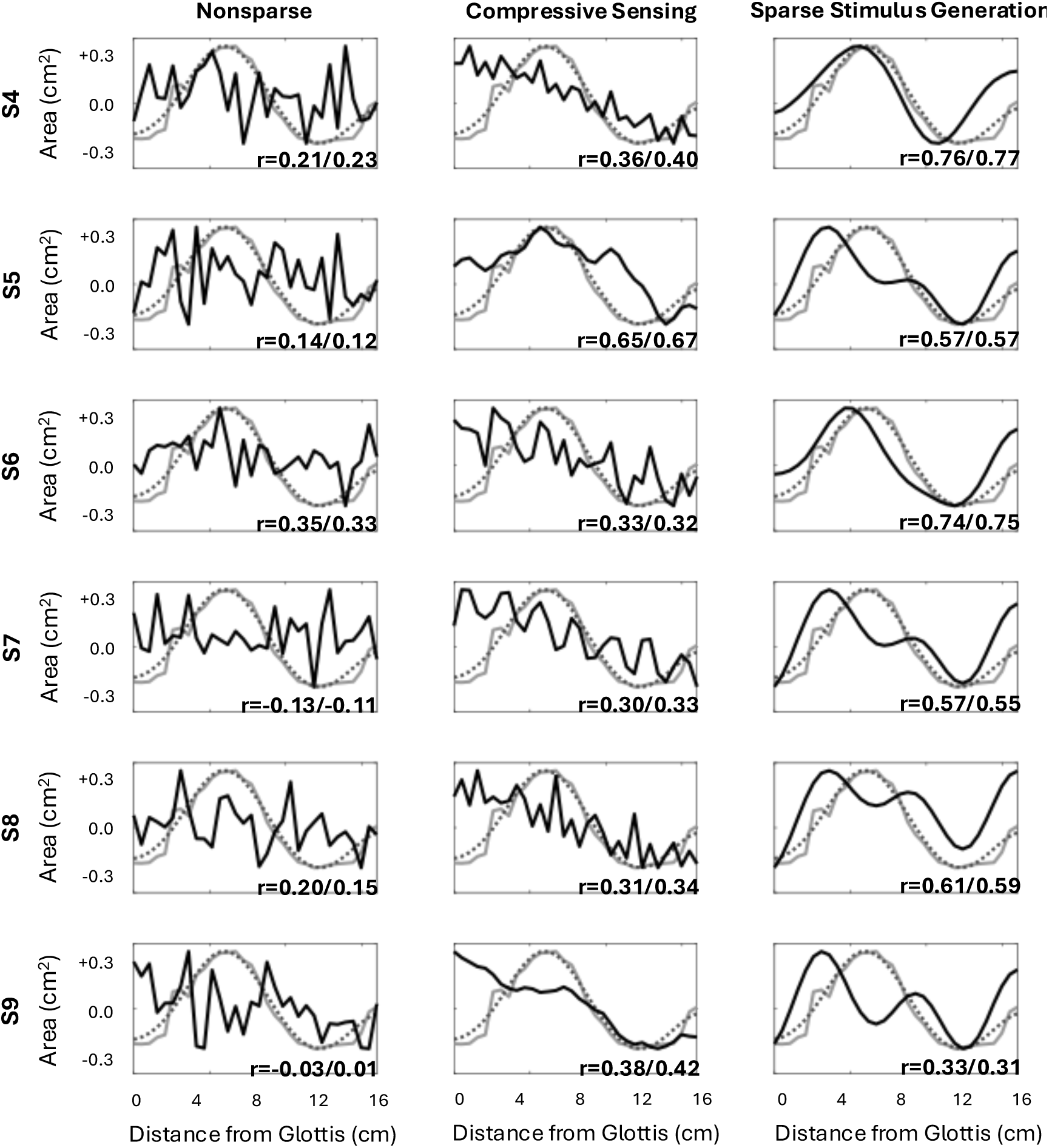
Additional reconstructions from human subjects. experiment 2 of the target vocal tract shape. Subfigures show reconstructions for a fixed level of sparsity (p=6) using the nonsparse, compressive sensing and sparse stimulus generation approaches (columns, left to right) and for a fixed number of trials (n=100) for each subject (rows). Estimation quality relative to the target signal b (gray, solid line), and a sparsified (p=6; gray, dotted line) version of b (as in Figure 1) are shown, respectively (target signal was resampled to m=32).

Subfigures show, across the columns, reconstructions for a fixed level of sparsity (p=6) and fixed number of trials (*n* = 200) using, from left to right: nonsparse stimulus generation with conventional reverse correlation reconstruction (Eq 4), nonsparse stimulus generation with compressive sensing reconstruction (Eq 5), and sparse stimulus generation with conventional reconstruction (Eq 4). Subfigure rows represent different subjects. Estimation quality in terms of Pearson’s *r* between the reconstruction and both the original target signal *b* (left of slash), as well as a sparsified (*p* = 6) version of *b* (right of slash).

In the results from human subjects experiment 1, sparse stimulus generation displayed consistently higher reconstruction quality than the conventional and compressive sensing approaches for any given number of trials, albeit with some variability in the magnitude of that improvement across subjects. The benefits of sparse stimulus generation decrease somewhat as the number of trials increased, with the reconstruction quality at 200 trials being approximately 115% above the nonsparse approach, as opposed to 350% when the number of trials was 40 (where the nonsparse reconstructions have almost no correlation with the target). Compressive sensing, while consistently worse than sparse stimulus generation, shows marginal improvement in reconstruction quality over conventional reverse correlation.

Table 1 shows the by-participant responses to questions about 1) similarity to the target and 2) confidence in positive responses from human subjects experiment 2. Values shown represent the mean response value across the four blocks presented for stimulus type. The bottom two rows of the table list the mean response value across subjects, as well as the standard deviations of the means (SD).

**Table 1:** Mean (n=4) Likert-scale responses for each subject to questions about similarity and confidence with respect to the sparse and nonsparse stimuli. Mean response value across subjects (N=24) and standard deviations of the means (SD) is also shown.

| subject | mean similarity (n=4) |  | mean confidence (n=4) |  |
| --- | --- | --- | --- | --- |
|  | nonsparse | sparse | nonsparse | sparse |
| S4 | 1.75 | 6.25 | 3.25 | 6.25 |
| S5 | 2.50 | 5.50 | 4.00 | 6.50 |
| S6 | 4.00 | 6.00 | 6.00 | 4.50 |
| S7 | 2.50 | 3.25 | 4.50 | 5.75 |
| S8 | 1.50 | 6.50 | 7.00 | 6.75 |
| S9 | 5.25 | 5.40 | 5.25 | 5.50 |
| Mean (N=24) | 2.92 | 5.48 | 5.00 | 5.88 |
| SD | 1.44 | 1.17 | 1.37 | 0.82 |

Participants found the sparsely generated stimuli (5.48 ± 1.17) to be much similar to the target sound than conventionally generated (nonsparse) stimuli (2.92 ± 1.44). This relationship held for every subject, with a mean Likert score difference of 2.56 across subjects (Cohen’s d = 1.95). Participants generally had more confidence in their positive response to the sparsely generated stimuli (5.88 ± 0.82) than the nonsparsely generated stimuli (5.00 ± 1.37), resulting in a mean Likert score difference of 0.88 across subjects (Cohen’s d = 0.78). Unlike the consistent responses for similarity, however, only 4/6 individual participants showed a pattern consistent with the overall group behavior.

## Discussion

Reverse Correlation is a widely-used technique for probing perceptual mechanisms. The technique involves presenting subjects with ambiguous stimuli across many trials. Preference scores are collected from the subject for each trial, for example a yes/no response with regard to whether the stimulus matches some latent perceptual representation. From the collected stimulus-response pairs, a reconstruction can be derived as an estimate of the representation that best explains subject responses given a linear comparison model of subject responses. It is often the case that the number of trials required for an adequate reconstruction is very large, which can be a major limitation to study design. Prior work from our group showed that the required number of trials can be lowered using a compressive sensing approach to reconstruction, in which it is assumed that the target representation may be sparsely represented in some basis. In the present work, we proposed a related approach in which the sparsity assumption in incorporated into the stimulus generation process, with the goal of lowering the number of required trials, while also making the stimuli less ambiguous for subjects. We applied this approach first in the context of a simulation study, with an idealized observer, in which the number of trials and the sparsity level were varied. Next, we applied this approach in in two behavioral studies with a total of nine three human participants, using a fixed number of trials and level of sparsity, in which the effect of the number of trials was assessed retrospectively.

Simulation study results reveal that, the sparse stimulus generation approach displays a higher reconstruction quality than the nonsparse, conventional reverse correlation approach for nearly any sparsity level and number of trials. These results are largely attributable to incorporating the assumption of sparsity of the target signal, in a way closely related to the sizeable body of results on compressive sensing, and owing to the close connections between sparse stimulus generation and compressive sensing. Notably, despite the close connections and shared assumptions of sparsity between sparse stimulus generation and compressive sensing, sparse stimulus generation consistently resulted in a substantially more accurate reconstruction than compressive sensing, although compressive sensing did produce higher quality reconstructions when compared to conventional reverse correlation reconstruction.

The benefits of sparse stimulus generation are most prominent when sparsity levels are high. When sparsity levels are very low, the performance of the sparse and nonsparse approaches converges. This effect is expected, as using a relatively large number of basis functions will produce stimuli that are effectively nonsparse, and which are likely to have properties similar to those of conventional reverse correlation stimuli.

Comparing simulation study results for sparse stimulus generation versus the compressive sensing approach, it is evident that sparse stimulus generation consistently displays a higher reconstruction quality for a wide range of sparsity levels and number of trials (although, at very low sparsity levels, compressive sensing outperforms both conventional reverse correlation and sparse stimulus generation for very small number of trials, sparse stimulus generation quickly starts to outperform compressive sensing as more trials are added). The poorer performance of compressive sensing when compared to sparse stimulus generation might also be expected, in that compressive sensing is solving a reconstruction problem which is inherently more difficult. Compressive sensing must simultaneously estimate not only the contribution of the basis functions, but must also determine which basis functions from amongst a larger set should be included in the sparse basis set. As noted above, however, the performance of compressive sensing appears much less sensitive to the number of trials when the value of *p* is large, which means that, when the level of sparsity is low, compressive sensing will require a relatively small number of trials to produce adequate reconstruction quality, but will require a large number of trials to produce excellent reconstruction quality. To our knowledge, this effect has not been previously reported in the literature on sparse reconstruction in reverse correlation (e.g., Mineault, Bartelme & Pack, 2009; Roop et al., 2024), and may merit further study.

Inspecting example reconstructions of the target signal (e.g., in Figure 3), it is noteworthy that the nonsparse stimulus approach is apparently able to reconstruct more detail of the target signal when compared to either the sparse stimulus approach or compressive sensing. While both sparse reconstructions are highly correlated with the target signal, the nonsparse reconstruction clearly preserves more of the fine details of the target signal. This effect is attributable, in the case of sparse stimulus generation, to the richer variation in the nonsparse stimuli, which more thoroughly explores the higher-dimensional space in with the original signal resides. In either case, this effect is a direct consequence of the assumption of sparsity, and demonstrates a potential drawbacks of that assumption, and the case which must be taken when applying it.

Results from the human subjects studies suggest that sparse stimulus generation produces much higher quality reconstructions than nonsparse stimulus generation, consistent with the simulation study results. Both sparse and nonsparse stimulus generation show a decreasing *rate* of improvement in accuracy with increasing number of trials, or *diminishing returns*, which has been noted many times in reverse correlation studies (Mineault, Barthelme & Pack, 2009; Compton et al., 2023; Roop et al., 2024). In contrast, the compressive sensing approach shows only marginal, if any, improvement in reconstruction quality as the number of trials increases, contrary to the simulation results. One possible explanation for this discrepancy may be the presence of perceptual noise in human participants, as opposed to the idealized observer assumed in the simulation study. However, Roop et al. (2024) showed that compressive sensing should be robust to this sort of noise. Another possible explanation might be that compressive sensing behaves worse than the nonsparse approach when sparsity levels are low (as shown in the simulation study). However, sparsity levels in the present human subject study were quite high. One additional explanation may be that the accuracy of the compressive sensing approach tends to exhibit a ceiling effect, such that it appears to converge to less-than-perfect accuracy, which cannot be easily remedied even with large numbers of trials (Roop et al. 2024). However, this performance ceiling is typically well above the 0.3-0.5 range observed in the present results. In short, the cause of the poor performance of compressive sensing in this study is unclear, and may merit further study.

Aside from the clear, quantifiable advantages of sparse stimulus generation in reconstruction quality, evidence from the subjective ratings in experiment 2 suggests that subjects find stimuli generated with the present approach to be more sensible and less perceptually ambiguous. All Subjects indicated strongly that the sparse stimuli were more similar to the target signal. Subjects also indicated, albeit with a smaller effect size and more variation between subjects, that they felt more confident when responding with a “yes” to the sparsely generated stimuli. These results suggest that the properties of sparsely generated stimuli may make reverse correlation easier and less fatiguing for participants.

Despite the apparent advantages of sparse stimulus generation in reverse correlation, care must be taken in applying the approach to specific studies with respect to the sparsity assumption and knowledge about the nature of sparsity in specific domains. As with compressive sensing, sparse stimulus generation is only appropriate in situations where sparsity of the target representations is a reasonable assumption. It has been argued here and in the broader reverse correlation literature that sparsity is a good assumption that is well-justified for many percepts of interest. Still, making the assumption of sparsity will necessarily bias results toward sparse estimates, even when the underlying representation is non-sparse.

Moreover, sparse stimulus generation has the added requirement that the domain of application must be one in which a well-justified sparse basis is known beforehand. Here, we have surveyed some examples of domains in which this is known, and with the sparsity assumption being widely accepted as an organizing principle in human perception, one might expect the number of application domains meeting this requirement of sparse stimulus generation to grow. At the same time, it is worth noting that bases for novel domains may be learned statistically using techniques like Principal Component Analysis and related, unsupervised machine learning methods. For example, reverse correlation could be used to gather a number of reconstructions in whatever quantity is feasible given the number of trials required for nonsparse methods. PCA could be applied to these reconstructions to infer an empirical basis for incorporation into sparse stimulus generation. Adopting sparse stimulus generation with such a basis for all subsequent data collection would then be substantially more efficient.

Sparse stimulus generation, like compressive sensing, can substantially decrease the number of trials required to reach high reconstruction quality in reverse correlation experiments by incorporating the assumption of sparsity. Provided that the somewhat stricter requirements of sparse stimulus generation can be satisfied, it can realize efficient improvements over and above what compressive sensing provides. Moreover, by incorporating sparsity into stimulus generation, at an earlier stage than reconstruction (as in compressive sensing), sparse stimulus generation holds promise for making the typically-ambiguous reverse correlation stimuli more sensible and easier to interpret for subjects. Both the efficiency gains and subjective enhancements hold promise for increasing sample sizes (in terms of the number of participants), easing participant attrition, and improving the overall quality of results in investigations of perceptual mechanisms.

## Acknowledgments

Primary support for this work was provided by a grant from the American Tinnitus Association. Student support was provided by the William F. McCall, Jr. ‘55 Summer Research Fund, as well as the JD Power Center at College of the Holy Cross.

